# Evaluation of Blood-Based Exosomes as Biomarkers for Aging-Related TDP-43 pathology

**DOI:** 10.1101/2022.06.03.487443

**Authors:** Charisse N. Winston, Sonal Sukreet, Haley Lynch, Virginia M.-Y. Lee, Donna M. Wilcock, Peter T. Nelson, Robert A. Rissman

## Abstract

**INTRODUCTION:** Limbic predominant age related TDP-43 encephalopathy (LATE) is a recently characterized brain disease that mimics Alzheimer’s disease (AD) clinically. To date, LATE is difficult to diagnose antemortem using clinical information or biomarkers. Recent studies suggest concentrations of exosomal protein cargo derived from neuronal and glial cells may serve as useful diagnostic biomarkers for AD and other neurodegenerative diseases.

**METHODS:** TDP-43 was evaluated in neuronal (NDE), astrocyte (ADE), and microglial derived exosomes (MDE). Exosome preparations were isolated from the plasma of research subjects with autopsy-confirmed diagnoses, including many with LATE. Quantified TDP-43 concentrations were compared to cohort that included healthy controls, mild cognitively impairment (MCI), and AD dementia with diagnoses other than LATE.

**RESULTS:** TDP-43 was significantly elevated in plasma ADEs derived from autopsy confirmed LATE-NC subjects, with or without comorbid AD pathology. Measurable levels of TDP-43 were also detected in exosome depleted plasma; however, TDP-43 levels were not significantly different between persons with and without eventual autopsy confirmed LATE-NC. No correlation was observed between exosomal TDP-43 levels with cognition-based variables, sex, and APOE carrier status.

**DISCUSSION:** Blood-based exosomes, specifically measuring TDP-43 accumulation in ADEs, may serve as a powerful diagnostic tool to rapidly identify subjects who are currently living with LATE-NC.

## INTRODUCTION

Abnormal accumulation of TAR DNA binding protein 43 (TDP-43) in its phosphorylated state has been associated with cognitive dysfunction and neurodegeneration in subjects who suffer from frontotemporal lobar degeneration (FTLD-TDP) [1, 2], amyotrophic lateral sclerosis (ALS) [1, 3], and Alzheimer’s disease (AD) [4]. Recently, limbic predominant age-related TDP43 encephalopathy (LATE) neuropathological change (NC) was characterized as distinct condition that is primarily marked by accumulation of TDP-43 proteinopathy in the hippocampus and entorhinal cortex of older adults [5]. Subjects with LATE-NC often have comorbid brain pathologies, including amyloid-β plaques and tau pathology, and display similar clinical manifestations to those who suffer with AD [5]. To date, LATE can only be diagnosed retrospectively (essentially impossible to diagnose with confidence in living subjects).

The need to identify neurodegenerative disease in the clinical setting, and at early and more treatable timepoints, has fueled research into blood-based biomarkers. Increased plasma concentrations of TDP-43 have been observed in subjects with FTLD [2]; however, its correlation with cognitive dysfunction and disease progression in AD is not well established [6]. Moreover, elevated plasma concentrations of TDP-43 in subjects with LATE has yet to be described.

Exosomes are nanosized extracellular vesicles (EVs) of endocytic origin that are released from various cell types, including neurons and glial cells [7]. Their diverse contents, including DNA, mRNA, and proteins, are thought to be reflective of the intracellular environment of the parent cell [7]. Secreted exosomes from neuronal cells are thought to mediate the propagation of TDP-43 in ALS and AD brains. Neuronal (NDE) and astrocyte (ADE) derived exosomes have been successfully isolated from the plasma of demented subjects and healthy controls [8–12]. While numerous studies suggest that protein cargo extracted from NDEs and ADEs may serve as useful diagnostic biomarkers for AD and other neurodegenerative diseases [8–12], very few studies have characterized the biomarker potential of protein cargo extracted from microglial derived exosomes (MDEs) [13, 14].

In the current study, TDP-43 accumulation was quantified in NDE, ADE, and MDE preparations isolated from the plasma of research subjects with autopsy-confirmed diagnoses, including many with LATE neuropathologic changes (LATE-NC(+)). Quantified TDP-43 concentrations were compared in a cohort that included healthy controls, mild cognitively impairment (MCI), and AD dementia with diagnoses other than LATE-NC (−).

## METHODS

### Baseline Characteristics of Subjects

Plasma samples from autopsy-validated LATE-NC positive and negative cases were acquired through the University of Kentucky Alzheimer’s Disease Research Center Biobank. Patient characterization details are listed in **Table 1**. LATE positive subjects had neuropathological changes associated with LATE-NC(+) and LATE-NC (−) subjects had no TDP-43 proteinopathy [5, 26]. Final clinical diagnoses were rendered at a consensus conference and varied between cognitively normal controls (CNC); subjects with an established diagnosis of mild cognitive impairment (MCI), Possible AD (AD), and subjects with Probable AD.

**Table 1.** Patient demographics for the 64 patient’s blood samples that were included in the study. Exosome preparations were enriched against neuronal (NDE), astrocyte (ADE), and microglial sources (MDE). Exosome cargo protein, TDP-43 was measured by human ELISA. Categorical variables are summarized using counts and percentages (%) with continuous variables summarized using means and standard deviations (SD).

| **Patient Demographic Data (n = 64)** |  |  |  |  |
| --- | --- | --- | --- | --- |
| **Characteristic** |  | **LATE – NC (-)**<br>**(N = 42)** | **LATE – NC (+)**<br>**(N = 22)** | **Combined**<br>**(N = 64)** |
| Age of Death, mean (SD) |  | 80 (9.6) | 84 (9.7) | 81 (9.8) |
| MMSE, mean (SD) |  | 22 (8.0) | 17 (11) | 21 (9.2) |
| Years of Education, mean (SD) |  | 16 (3.6) | 18 (2.9) | 17 (3.4) |
| **Last Clinical Index** | Control | 13 (31%) | 3 (13.6%) | 16 (25%) |
|  | MCI or Impaired | 9 (21.4%) | 3 (13.6%) | 12 (18.75%) |
|  | Demented (AD) | 20 (47.6%) | 16 (72.7%) | 36 (56.25%) |
| **Sex** | Female | 21 (50.0%) | 9 (40.9%) | 30 (46.9%) |
|  | Male | 21 (50.0%) | 13 (59.1%) | 34 (53.1%) |
| **APOEε4** | Negative | 24 (57.1%) | 10 (45.5%) | 35 (54.7%) |
|  | Positive | 17 (40.5%) | 11 (50%) | 28 (43.8%) |
|  | (Missing) | 1 (2.4%) | 1 (4.5%) | 2 (1.5%) |
| **Race** | White | 40 (95.2%) | 21 (95.5%) | 61 (95.3%) |
|  | Black or African American | 2 (4.8%) | 1 (4.5%) | 3 (4.7%) |
|  | Hispanics | 0 (0.0%) | 0 (0.0%) | 0 (0.0%) |
|  | Asian | 0 (0.0%) | 0 (0.0%) | 0 (0.0%) |
|  | American Indian or Alaska Native | 0 (0.0%) | 0 (0.0%) | 0 (0.0%) |

### Isolation and neuronal enrichment of blood-based exosomes derived neuronal (NDEs), astrocyte (ADEs) and, microglial (MDEs) sources

Exosome isolation was conducted as previously described [1]. Briefly, 500 μL of human plasma were incubated with 5 μL purified thrombin (System Biosciences, Inc.; Catalog # TMEXO-1) at room temperature for 5 min. After centrifugation at 10,000 rmp for 5 min, supernatants were incubated with 120 μL ExoQuick Exosome Precipitation solution (System Biosciences, Inc.; Catalog # EXOQ5TM-1) for 30 min at 4°C. Resultant suspensions were centrifuged at 1,500 × g for 1 h at 4°C. Supernatant was collected and the resultant pellet was suspended in 350 μL of 1× phosphate buffer saline (PBS) (diluted from 10× PBS; Thermo Fisher Scientific; Catalog# AM9625) with Halt protease and phosphatase inhibitor cocktail EDTA-free (Thermo Fisher Scientific; Catalog # 78443) and stored at −80°C until immunochemical enrichment of exosomes from both neural and astrocytic sources.

Neuronal, astrocyte, and microglial enrichment was conducted per manufacturer’s instructions (System Biosciences, Inc.; Catalog # CSFLOWBASICA-1). Briefly, 45 μL of 9.1 μm, streptavidin magnetic Exo-Flow beads (System Biosciences, Inc.; Catalog # CSFLOWBASICA-1) were incubated with 100 ng/μL of mouse anti-human CD171 (L1CAM, neural adhesion protein) biotinylated antibody (clone 5G3, eBioscience/Thermo Fisher Scientific; Catalog # 13-1719-82); mouse anti-human GLAST (ACSA-1) biotinylated antibody (Miltenyi Biotec, Inc., Auburn, CA, United States; Catalog # 130-118-984; or Purified anti-TMEM119 (Extracellular) Antibody (Biolegen, Catalog # 853302) for 2 h on ice, with gently flicking every 30 min to mix. Anti-TMEM119 was biotinylated prior to exosome enrichment (Biotinylation Kit / Biotin Conjugation Kit (Fast, Type A) - Lightning-LinK, Catalog # ab201795). Bead-antibody (Ab) complex was washed three times in Bead Wash Buffer (Systems Biosciences, Inc.; CSFLOWBASICA-1) using a magnetic stand. Bead-Ab complex was suspended with 400 μL of Bead Wash Buffer and 100 μL of total exosome suspensions rotating overnight at 4°C. Bead-Ab-exosome (BAE) complex was washed three times with Bead Wash Buffer then suspended in 240 μL of Exosome Stain Buffer and 10 μL of Exo-FITC Exosome FACS stain (Systems Biosciences, Inc.; Catalog # CSFLOWBASICA-1) for 2 h on ice, with gently flicking to mix. BAE-FITC complex was washed Three times in Bead Wash Buffer then suspended in 300 μL of Bead Wash Buffer prior to loading into BD FACS Aria II for sorting. Flow-sorted, BAE-FITC complexes were incubated with 350 μL of Exosome elution buffer (System Biosciences, Inc.; Catalog # CSFLOWBASICA-1) at 25°C for 30 min. Finally, supernatant containing eluted exosomes were incubated with 1 μL of Exo-FlowIP clearing reagent (System Biosciences, Inc.; Catalog # EXOFLOW32A) at 37°C for 30 min then stored at −80°C.

Nanoparticle Tracking Analysis (NTA) was used to characterize L1-CAM-positive (NDEs), GLAST-positive (ADEs), and TMEM119-positive (MDEs) based on size distribution. Exosome preparations were confirmed by tetraspanning exosome marker CD81 (Cusabio, American Research Products, Waltham, MA, United States; Catalog # CSB-EL004960HU).

### TD-43 Assay

Protein concentrations for all exosome preparations were determined using a bicinchoninic acid (BCA) Protein Assay kit (Pierce Biotechnology). Exosome cargo proteins were quantified by commercially available human-specific ELISAs kits for Human TAR DNA-binding protein 43 (TARDBP/TDP43) (CSB-E17007h; Cusabio; Waltham, MA); Human TDP-43 DuoSET ELISA kit DuoSet Ancillary Reagent Kit 2 (#DY008, R&D Systems, Minneapolis, MN, USA), and tetraspanin exosome marker CD81 (CSB-EL004960HU; Cusabio; Waltham, MA) according to suppliers’ directions.

Briefly, 96 well plates were coated overnight with capture Ab Human Recombinant TDP-43 (mAb 5044.48.42(7-12-16), IgG1, aa 261 – 393, gift from Dr. John Q. Trojanowski, UPENN), washed 3× in 1× wash buffer (895003, R&D Systems), blocked in 1% BSA for 1 hour at room temperature (RT). Coated plates were washed 3× in 1× wash buffer, air dried overnight at 37°C and then stored at 4°C in sealed plastic covers prior to use. Coated plates are incubated with 100 ul exosome preparations and standards overnight at 4°C followed by Human TDP-43 - biotin detection antibody (Recombinant Anti-TDP43 antibody (ab255922) + Biotinylation Kit/Biotin Conjugation Kit (Type A) (ab201795)) for 2 hours at RT. Signal detection of TDP-43 protein concentration was quantified by Streptavidin-HRP and then measured at an optical density of 450 nm using a microplate reader (Bio-Rad, Hercules, California, USA). Absorbance values were converted into concentrations using a standard curve of TDP-43 in the range of 0 to 20 ng mL – 1. The mean value for all determinations of CD81 in each assay group was set at 1.00, and the relative values for each sample were used to normalize their recovery.

### Statistical analyses

The statistical significance of differences between means for cross-sectional patient groups and their respective control group was determined by an unpaired, non-parametric Mann–Whitney t-test or a one-way ANOVA with Newman-Keuls Multiple Comparison post hoc test (Prism 9; GraphPad Software, La Jolla, CA, United States).

## RESULTS

Sixty-four subjects were included in the study: twenty-two autopsy confirmed LATE-NC (+) (mean age at death, 84 ± 9.7 years) and forty-two subjects without LATE-NC (−) diagnosis (mean age at death, 80 ± 9.6 years). The percentage of *APOE* ε4 carriers (50% vs. 40.5%) and the percentage of males vs. females (males, 46.9%, females, 53.1%) were similar across both groups. The LATE-NC (+) group contained a significantly higher number of autopsy-confirmed AD neuropathologic changes (ADNC) [15] (72.7% vs. 47.6%) with a lower MMSE score (17 vs. 22) as compared to the LATE-NC (−) group.

Plasma NDEs, ADEs, and MDEs were characterized by FACs sorting [8], nanoparticle tracking analysis (NTA) [9], and human specific ELISAs. The significant degree in flow separation from the non-exosome negative control suggested a successful enrichment of exosome preparations against neuronal, astrocyte, and microglial sources (**Fig.1A**). Extracted plasma exosomes were similar in size to previously published studies (**Fig. 1B**) [9]. Assessments of the exosome membrane marker CD81 confirmed resultant exosome preparations that were derived from LATE-NC (+) and LATE-NC (−) samples. Concentrations of CD81 were not statistically different between NDE, ADE, and MDE preparations (**Fig. 1C**).

**Figure 1.**
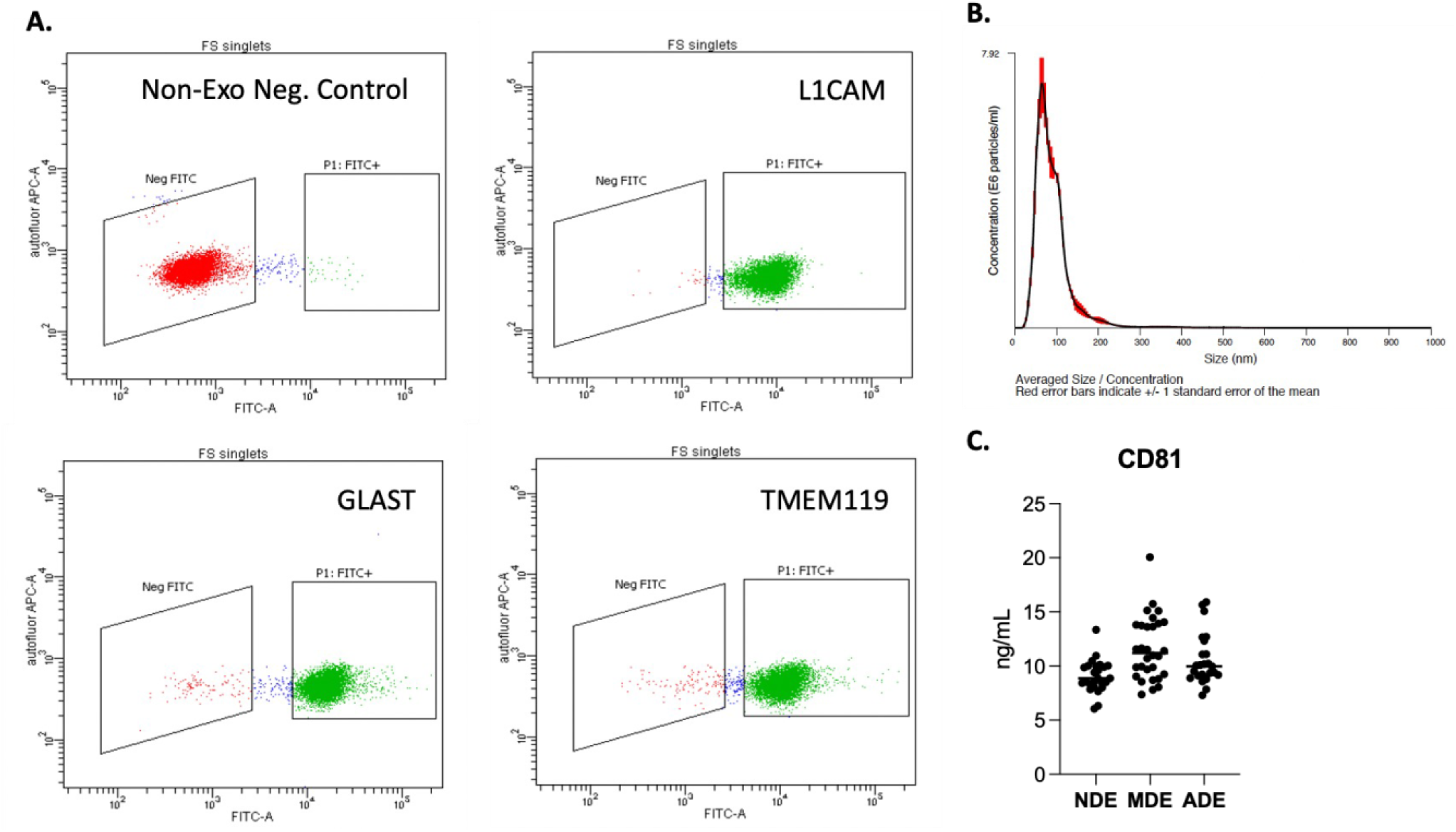
Fluorescent activated cell sorting (FACS) enrichment of plasma NDEs, ADEs, and MDEs from LATE-NC subjects. Representative FACS plot for non-exosome, negative control (red) and BAE – FITC complexes generated from exosomes (green) isolated a LATE-NC subject and enriched against anti-human CD171 biotin (L1CAM), anti-GLAST, and anti-TMEM119 antibody. **(A)** Representative plot of size/concentration determined by nanoparticle tracking analysis (NTA) for extracted plasma exosomes from a LATE-NC subject **(B)**. Plasma concentrations of exosome marker, CD81 as measured by hu-specific ELISA. CD81 was not statistically different between the three groups **(C)**.

A direct comparison between an in-house, sandwich-based ELISA and the commercially available Cusabio kit was conducted in this study. TDP-43 protein concentration was measured in NDE, ADE, and MDE preparations from all samples. Across both ELISA platforms, NDE concentrations of TDP-43 were not significantly different between LATE-NC (+) and LATE-NC (−) subjects (**Fig. 2A – B**). ADE concentrations of TDP-43 were significantly increased in LATE-NC (+) as compared to LATE-NC (−) subjects (**Fig. 2C – D**). MDE concentrations of TDP-43 were significantly increased in LATE - NC (+), as measured by our in-house ELISA (**Fig. 2E**), while MDE concentrations of TDP-43 were significantly decreased in LATE-NC (+), as measured by the Cusabio kit (**Fig. 2F**). In exosome depleted plasma, measurable levels of TDP-43 concentrations were not significantly different between LATE-NC (+) and LATE-NC (−) subjects. These findings were consistent across both ELISA platforms. (**Fig. 2G – H**). In sum, significantly higher levels of TDP-43 protein were detected by our in-house ELISA kit as compared to the Cusabio kit in the NDE, ADE, and MDE exosome preparations and in the exosome depleted plasma (1.56 ± 0.078 ng/mL vs. 0.66 ± 0.089 ng/mL, mean ± SEM, P < 0.001, **Fig. 2G – H**).

**Figure 2.**
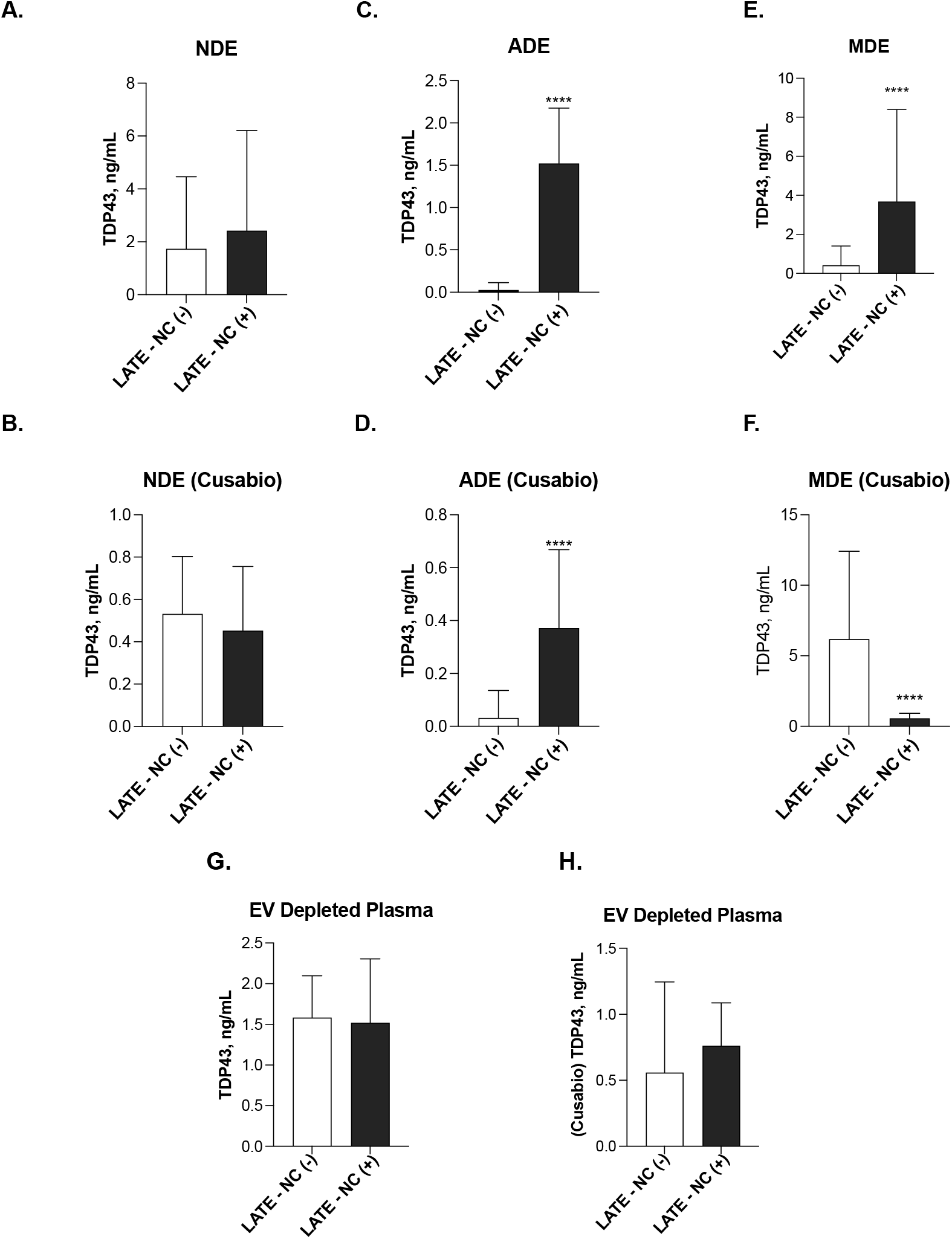
TDP-43 is elevated in ADEs extracted from subjects with an autopsy-confirmed diagnosis of LATE. Plasma concentrations of TDP-43 were detected in NDEs, ADEs, and MDEs **(A -F)** and in exosome depleted plasma **(G – H)** as measured by our in-house ELISA and Cusabio kit. NDE concentrations TDP-43 was not significantly different between LATE-NC (+) and LATE-NC (−) subjects (**A – B)**. ADE concentrations of TDP-43 were significantly increased in LATE-NC (+) as compared to LATE-NC (−) subjects (**C – D**). MDE concentrations of TDP-43 were significantly increased in LATE - NC (+), as measured by our in-house ELISA (**Fig. 2E**), while MDE concentrations of TDP-43 were significantly decreased in LATE-NC (+), as measured by the Cusabio kit. Exosome depleted plasma concentrations of TDP-43 were not significantly different between LATE-NC (+) and LATE-NC (−) subjects.

Correlations between TDP-43 accumulation in NDEs, ADEs, and MDEs against cognition-based variables including MMSE score and Braak NFT staging; sex; and *APOEε4* carrier status were established to compare results between LATE-NC (+) and LATE-NC (−) subjects. As determined by our in-house platform, no correlation was observed with TDP-43 accumulation in NDEs, ADEs, and MDEs with a lower MMSE score (**Fig. 3A – C**) or Braak NFT stage (**Fig. 3D – F**). Moreover, there was no significant difference in NDE, ADE, and MDE concentrations of TDP-43 between females vs. males (**Figures 3G – I**) or between *APOEε4* carriers and non-carriers (**Fig. 3J – L**). Similarly, no correlation was observed with TDP-43 accumulation in NDEs, ADEs, and MDEs as measured by the Cusabio kit (**Fig. 4A – L**).

**Figure 3.**
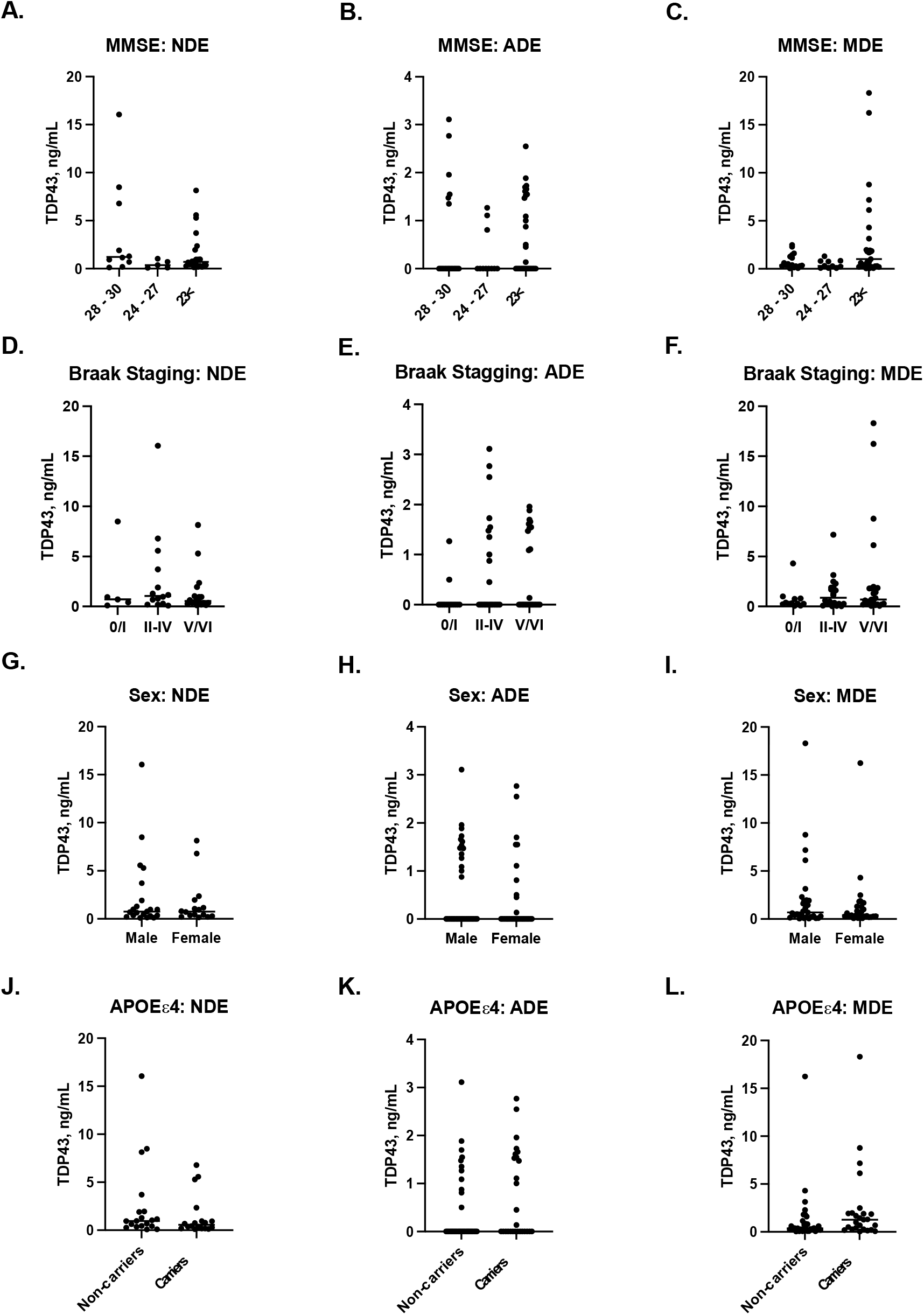
Correlations between TDP-43 accumulation in NDEs, ADEs, and MDEs against cognition-based variables including MMSE score and Braak NFT staging; sex; and *APOEε4* carrier status as measured by our in-house ELISA. No correlation was observed with TDP-43 accumulation in NDEs, ADEs, and MDEs with a lower MMSE score (**A – C**) or Braak NFT stage (**D – F**). NDE, ADE, and MDE concentrations of TDP-43 were not significantly different between females vs. males (**G – I**) or between *APOEε4* carriers and non-carriers (**J – L**).

**Figure 4.**
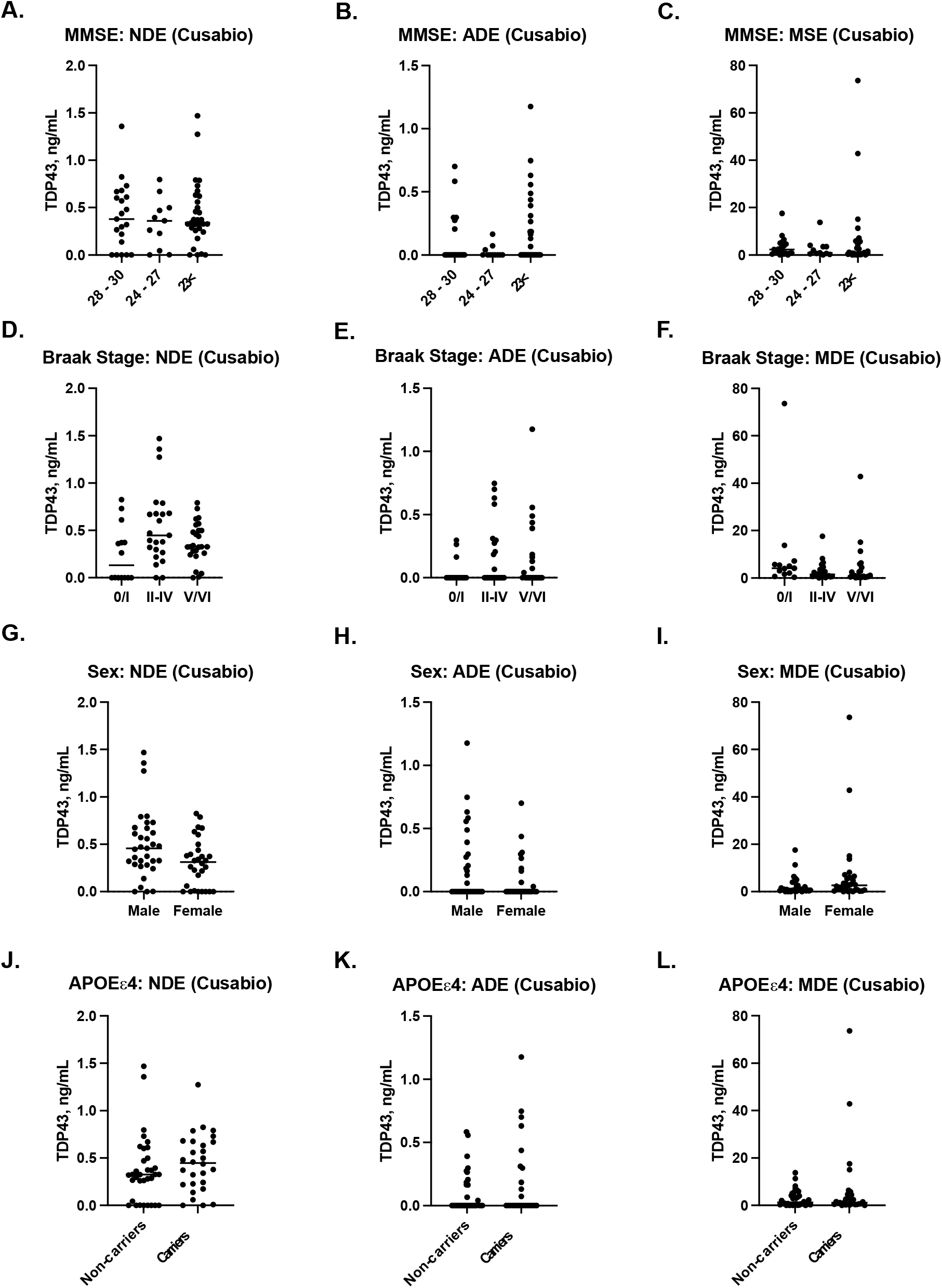
Correlations between TDP-43 accumulation in NDEs, ADEs, and MDEs against cognition-based variables including MMSE score and Braak NFT staging; sex; and *APOEε4* carrier status as measured by our Cusabio kit. No correlation was observed with TDP-43 accumulation in NDEs, ADEs, and MDEs with a lower MMSE score (**A – C**) or Braak NFT stage (**D – F**). NDE, ADE, and MDE concentrations of TDP-43 were not significantly different in females vs. males (**G – I**) or between *APOEε4* carriers and non-carriers (**J – L**).

## DISCUSSION

In the current study, we generated a sensitive, sandwich-based ELISA that can detect TDP-43 in NDE, ADE, and MDE preparations isolated from human plasma, and we compared between LATE-NC (+) and LATE-NC (−) subjects. We determined that ADE cargo levels contain significantly elevated levels of TDP-43 in subjects with an autopsy-confirmed diagnosis of LATE-NC. These data suggest that plasma ADEs may serve as useful diagnostic biomarker to identify subjects with LATE-NC. To our knowledge, this is the first study to report these findings.

Measurable levels of TDP-43 was also detected in exosome depleted plasma; however, TDP-43 levels were not significantly different between persons with and without eventual autopsy confirmed LATE-NC. Our results indicate that there is a substantial diagnostic advantage to measuring exosomal levels of TDP-43. There is accumulating evidence suggesting TDP-43 induces proinflammatory cytokines release from astrocytes and dysregulates astroglial metabolic support of neurons in ALS [16] and FTD [17]. However, the biological rationale for these molecular underpinnings are currently not fully understood and await future studies. In the meantime, without understanding the underlying biology, the ADE TDP-43 may still provide the basis for a useful clinical biomarker. We note that our “in-house” ELISA detected higher levels of exosomal TDP-43 as compared to the Cusabio ELISA, and consider it likely that differences are due to the specificity of the TDP-43 detection antibodies that were used in both platforms [18].

A limitation of the current student is the sample size. The small sample size may partly explain our inability to identify a correlation between exosomal TDP-43 levels with cognition-based variables, sex, and APOE carrier status. However, we do note that there was very robust association between ADE TDP-43 levels and LATE-NC status even with the sample size that was used, and despite the inclusion of a broad spectrum of dementia diagnosis controls. Prior studies have found an association between *APOE* genotype with AD [19] LATE-NC [20]. Zhang et al recently reported that TDP-43 is elevated in plasma NDEs of subjects with AD yet they did not observe a correlation with TDP-43, APOE genotype and a collection of neuropsychiatric symptoms [18]. Similarly, we did not observe an association between NDE levels and TDP-43 in the present study. The lack of ethnoracial diversity in the current study is another major limitation. In our sample, 95% of subjects were identified as non-Hispanic Caucasian. Hispanics/Latinos and African Americans/blacks have a higher prevalence of AD as compared to non-Hispanic whites [21]. Early studies indicate that LATE-NC occurs with similar prevalence in African Americans/blacks and whites [22]. Recent observational studies have reported the impact of race on AD biomarker performance [23-25] but more work is urgently required in the clinical, pathological, genetic, and biomarker domains with the goals of better serving broad populations [22]. Ultimately, our study must be validated in a larger and more diverse patient cohort.

In conclusion, LATE is an under-recognized condition with a large impact on public health for which, to date, the lack of a clinical biomarker is a glaring unmet need [5]. In the current study, we found a dramatic difference in ADE levels of TDP-43 in LATE-NC, suggesting that there is a unique interaction with TDP-43 and astrocytes that also requires further investigation. Blood-based exosomes, specifically measuring TDP-43 accumulation in ADEs, may serve as a powerful diagnostic tool to rapidly identify subjects who are currently living with LATE-NC.

## Acknowledgements

This work was supported by NIH grants to RR (AG0518440, AG051848, AG058533, AG062429), K99/R00 AG070390 to CNW.

